# Robust Neural Decoding with low-density EEG

**DOI:** 10.1101/2025.07.07.663494

**Authors:** Ling Huang, Manuel Varlet, Tijl Grootswagers

## Abstract

High-density Electroencephalography (EEG) recording enhances spatial resolution for neural signal decoding, yet the relationship between electrode density and decoding performance remains unclear. To address this, we systematically investigated decoding accuracy across electrode configurations of varying densities (16, 32, 64, 96, and 128 electrodes) using visual grating stimuli characterized by orientation, contrast, spatial frequency, and color. As expected, decoding accuracy increased with electrode density. Remarkably, however, reliable above-chance decoding was still achieved with as few as 16 electrodes, highlighting the robustness of decodable neural signals. To test the generalization of these results to more complex natural stimuli, we conducted a similar analysis with a diverse set of naturalistic images categorizable into living/non-living and moving/non-moving. The results consistently showed that effective decoding persists even with a 16-electrode configuration, showing robust decoding efficacy even for complex naturalistic stimuli. These findings demonstrate both the benefits of higher-density EEG and the robustness of neural decoding under sparse spatial sampling, providing new insights into how efficiently and broadly neural signals can be decoded.

## 1. Introduction

Electroencephalography (EEG) is a widely used non-invasive technique with high temporal resolution that has become a valuable tool for neuroimaging studies^1^. In particular, it is extensively used in visual studies to investigate cognitive processes such as object recognition, spatial attention, working memory, and motion perception^2–7^. Numerous studies using diverse visual stimuli, from simple gratings to natural scenes, suggest that EEG signals encode rich information about both low- and high-level visual representations^4–7^.

Despite these versatile neural representations across stimulus types, the extent to which the number of electrodes affects decoding performance remains controversial^8–10^. With the advancement of EEG technology, the number of electrodes used in recording has steadily increased, leading to the development of high-density EEG systems. Generally, these systems offer enhanced spatial resolution and improved neural signal decoding capabilities^10–12^. However, this does not mean that lower-density EEG cannot support meaningful decoding. Recent studies have suggested that image classification, image reconstruction, and functional connectivity analysis may still be feasible using low-density EEG, challenging the assumption that more electrodes necessarily yield accurate decoding performance^8,9,13^.

From a neurophysiological perspective, this possibility is supported by the biophysical properties of EEG. Although EEG signals originate from anatomically localized sources, such as early visual areas in the occipital cortex involved in encoding low-level features^14^, the resulting scalp potentials are shaped by volume conduction, which causes neural activity to spread through the brain and skull. This process produces spatially widespread but locally enhanced scalp fields^15–18^. As a result, sparse but broadly distributed EEG montages, when designed to preserve full-head coverage, may still capture the key components of multivariate signal patterns, allowing even low-density electrode configurations to retain sufficient information for reliable decoding^8,9,13,19^.

Moreover, deploying high-density EEG is associated with several limitations, including high costs, extensive setup time, lack of portability, and substantial computational demands^8,9,13^. In contrast, low-density EEG has been shown to be effective in brain-computer interface (BCI) applications, particularly among individuals with motor disabilities^20^. In clinical practice, especially in epileptology, low-density EEG is routinely used, with 9 to 21 electrodes commonly employed for diagnostic purposes^21,22^. These observations raise a critical question: can comparable decoding performance be achieved with a reduced number of electrodes, thereby making EEG-based decoding more practical and widely accessible?

To address this issue, we systematically investigated the impact of electrode density on visual decoding performance. Using a publicly available EEG dataset^23^ featuring visual grating stimuli varying in orientation, contrast, spatial frequency, and color, we examined decoding accuracy across electrode configurations of different densities (16, 32, 64, 96, and 128 electrodes). As expected, decoding accuracy increased with electrode density. Remarkably, however, reliable above-chance decoding was still achieved with as few as 16 electrodes, highlighting the robustness of decodable neural signals. To assess the generalizability of these results, we extended our analysis to a diverse set of natural images from another publicly available EEG dataset^24^. This dataset included visual image categories such as living animals (whales), living plants (flowers), non-living moving artificial objects (trains), non-living moving natural objects (waterfalls), non-living still artificial objects (cups), and non-living still natural objects (rocks). Using these natural stimuli, we again found that decoding performance remained robust even with as few as 16 electrodes, suggesting that high-level visual information can also be reliably captured with low-density EEG. The consistency of decoding performance across these varied stimulus reveal that while higher-density EEG improves decoding performance, reliable neural decoding can still be achieved with sparse electrode configurations, underscoring the efficiency and generalizability of neural signal representations.

## 2. Methods

This study utilized publicly available datasets from OpenNeuro: Experiment 1 (https://doi.org/10.18112/openneuro.ds004357.v1.0.1) and Experiment 2 (https://doi.org/10.18112/openneuro.ds003885.v1.0.7). In Experiment 1, participants were presented with oriented grating stimuli, shown for one frame (16.67 ms) in sequences at 6.67 Hz (133 ms inter-stimulus interval, ISI; 150 ms stimulus onset asynchrony, SOA) or 20.00 Hz (33 ms ISI; 50 ms SOA), and were asked to fixate on a black bullseye and detect when the fixation bullseye changed to a filled circle target^23^. In Experiment 2, participants performed a categorization task with natural stimuli, classifying images based on the criterion “whether they are alive or not,” as well as a passive viewing task, during which they viewed stimuli presented in rapid streams^24^. To investigate how electrode density influences decoding performance, we reanalyzed the 128-channel electroencephalography (EEG) data from both experiments using different standard electrode density configurations (16, 32, 64, 96, and 128 electrodes; Fig. 1; Supplementary Table 1). We applied the exact analysis pipelines from the original studies to the reduced configurations, and summarize these analysis pipelines below.

**Fig. 1.**
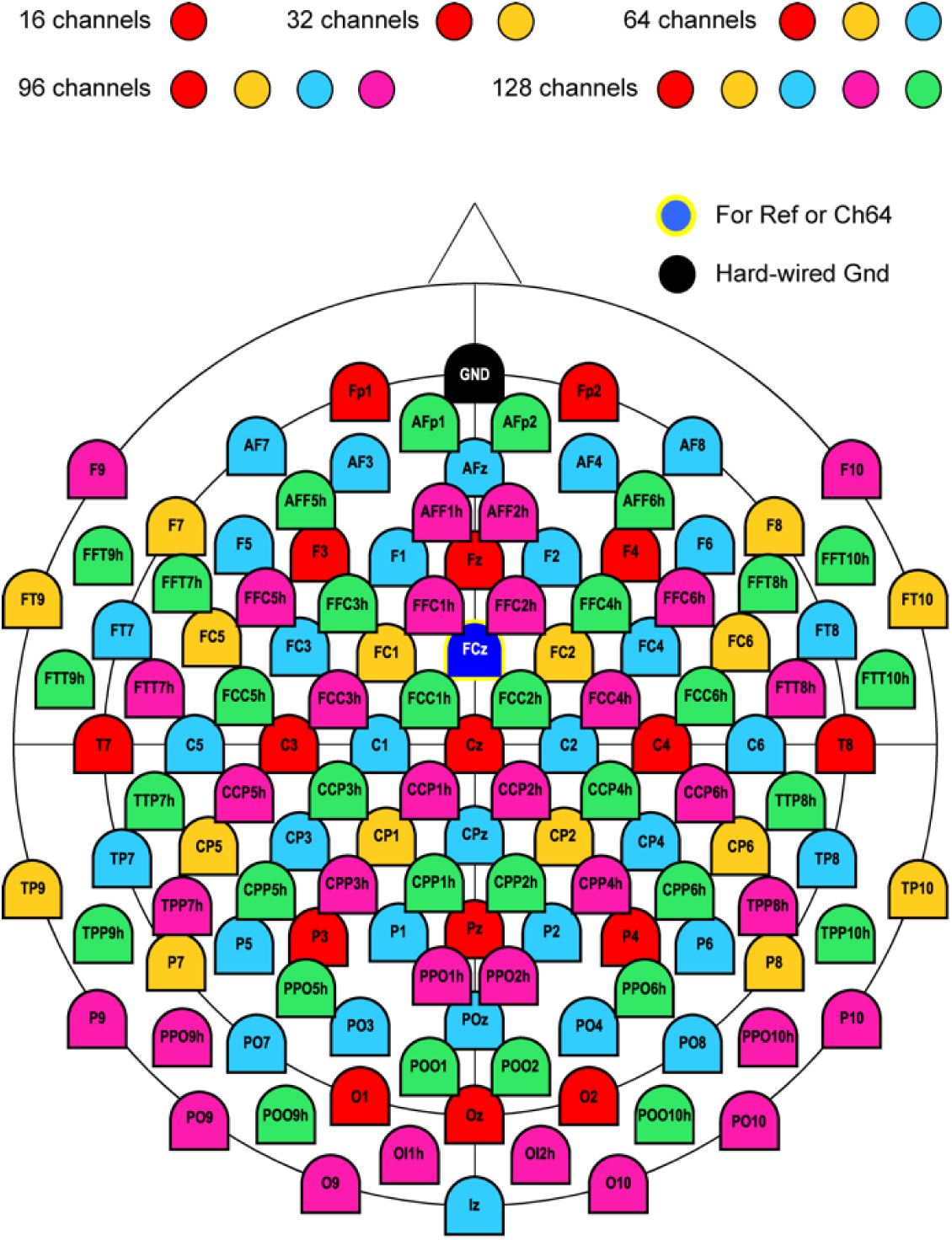
Electrode channel layouts for each electrode density configuration. Schematic layout of the 128-channel EEG montage with subsets used for different electrode density configuration. Colored electrodes indicate the five density levels: 16 channels (red), 32 channels (red and yellow), 64 channels (red, yellow and blue), 96 channels (red, yellow, blue and magenta), and 128 channels (red, yellow, blue, magenta and cyan). Subsets were selected to follow standard low-density EEG configurations and were evenly distributed across the scalp. The black electrode indicates the hard-wired ground (Gnd), and the blue with a yellow outline electrode indicates the reference channel (Ch64).

### 2.1 Participants

The analyses in this study were based on publicly available EEG datasets originally collected at the University of Sydney. The original experiments were conducted under University of Sydney ethics committee approvals, and informed written consent was obtained from all participants. Experiment 1^23^ involved 16 participants (11 females and 5 males, age range: 18–27 years), while Experiment 2^24^ included 24 participants (15 females and 9 males, age range: 18–26 years).

### 2.2 Apparatus

In Experiment 1, stimuli were presented centrally with a visual angle of 6.5°, while in Experiment 2, the visual angle was approximately 5°. In both experiments, EEG data were recorded while participants viewed experimental stimuli presented on a monitor with a 60 Hz refresh rate. Recordings were acquired at a sampling rate of 1000 Hz using a 128-channel BrainVision ActiCap system (Brain Products GmbH). Electrode placement followed standard scalp configuration guidelines, using the international five percent system in both experiments^25^. EEG signals were referenced online to FCz.

### 2.3 Stimuli

Both original experiments utilized well-documented stimulus sets. As described in the original study, the stimuli in Experiment 1^23^ consisted of 256 oriented sinusoidal gratings generated using the GratingStim function in PsychoPy^26^. Each stimulus subtended 6.5° of visual angle and varied systematically along four visual feature dimensions: color, spatial frequency, contrast, and orientation (Fig. 2A). Four levels were defined for each feature dimension. The four colors were RGB values ([66, 10, 104], [147, 38, 103], [221, 81, 58], [252, 165, 10]) that were approximately equidistant in color space, and each was paired with its complementary color. Orientations (22.5°, 67.5°, 112.5°, 157.5°) were evenly spaced circularly, while spatial frequencies (2.17, 1.58, 0.98, 0.39 cycles/°) and contrast values (0.9, 0.7, 0.5, 0.3) were linearly spaced. The resulting stimulus space included all 256 possible combinations. To prevent potential confounds related to phase, the phase of each grating was randomly varied on each presentation.

**Fig. 2.**
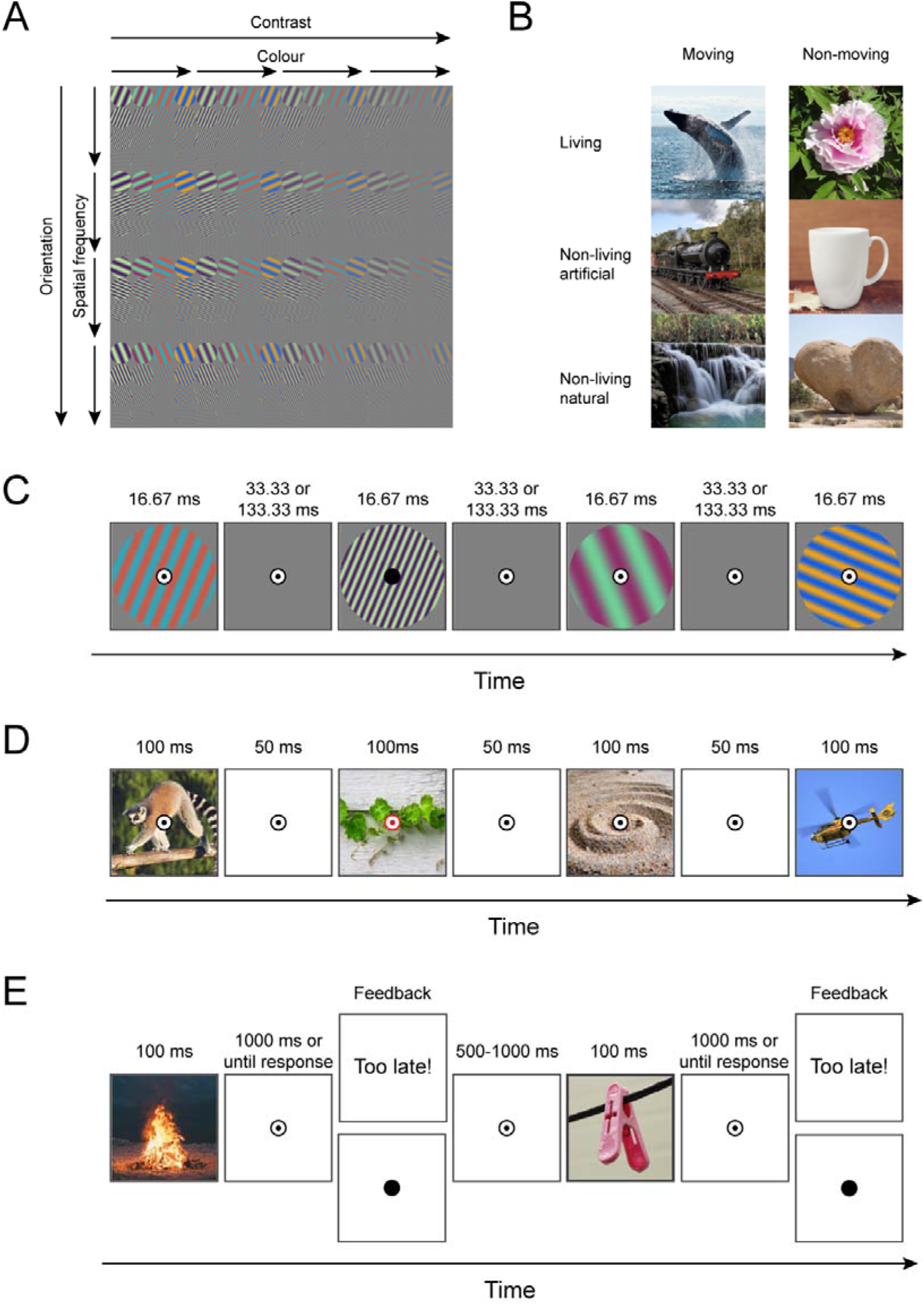
Experimental paradigm for Experiments 1 and 2 with EEG. Sample stimuli are shown in (A) for Experiment 1 and (B) for Experiment 2. In Experiment 1 (C), visual grating stimuli were presented for one frame (16.67 ms) at 6.67 Hz (133 ms ISI; 150 ms SOA) or 20.00 Hz (33 ms ISI; 50 ms SOA). Participants were asked to detect when the fixation bullseye changed to a filled circle target and respond using a button press. In Experiment 2, during passive viewing trials (D), participants viewed a rapid stream of natural images and responded by pressing a button when the fixation spot changed to red. During categorization trials (E), participants categorized images based on whether each depicted something alive or not alive. Note that all images are magnified here for clarity; in the actual presentation, they occupied a smaller proportion of the screen. Panels (A) and (C) are adapted from Grootswagers et al.^23^, while Panels (B), (D), and (E) are adapted from Shatek et al. ^24^.

The stimuli in Experiment 2^24^, as described in the original study, comprised 400 naturalistic color images sourced from publicly available image databases (www.pixabay.com and www.pexels.com) under Creative Commons 0 licenses. All images were manually processed by the original authors using GIMP (v2.10.14)^27^ to blur identifiable text (e.g., brand names) and were subsequently cropped and resized to approximately 5° of visual angle. The images were organized into six semantic categories based on Goldberg and Thompson-Schill^28^: animals (bee, cat, dog, dolphin, eagle, horse, lemur, pigeon, tiger, whale) and plants (cactus, clover, fern, flower, grass, lemon tree, moss, palm tree, tree, vine), which each included 10 objects. For all other categories, still artificial things (bench, clothes peg, headphones, lock, mug), still natural things (cliff, crystal, rock, sand, shell), moving artificial things (boat, bus, car, helicopter, train), and moving natural things (fire, hot spring, river, waterfall, waves), there were 5 objects. Within each category (e.g., cat, bench), there were 10 different images (e.g., cat1, cat2, … cat10). For objects capable of movement, the corresponding images depicted dynamic scenes (e.g., dolphins leaping out of the water, flowing waterfalls; see Fig. 2B), although all stimuli were presented as static images. Comprehensive information is provided in Shatek et al.^24^.

### 2.4 EEG experiment procedure

Neural responses were collected using electroencephalography (EEG) while participants viewed experimental stimuli and performed specific tasks, following protocols from the original studies^23,24^.

#### 2.4.1 Experiment 1

Participants viewed sequences of 256 grating stimuli, each presented for 16.67 ms at either 6.67 Hz (150 ms SOA) or 20.00 Hz (50 ms SOA). Each sequence consisted of all 256 stimuli presented in random order, and 80 sequences were shown in total, with each stimulus repeated 40 times per frequency. A fixation bullseye appeared one second before each sequence and remained superimposed throughout. Participants pressed a button when the bullseye briefly changed to a filled circle, which occurred 2 –4 times per sequence (Fig. 2C).

#### 2.4.2 Experiment 2

Participants completed eight blocks alternating between a categorization task and a passive viewing task. For the current reanalysis, only the “alive” category task (2 blocks) and the passive viewing task (4 blocks) were included. To balance trial numbers across the six natural stimulus categories: living animals (whales), living plants (flowers), non-living moving artificial objects (trains), non-living moving natural objects (waterfalls), non-living still artificial objects (cups), and non-living still natural objects (rocks), half of the animal and plant trials were randomly selected.

For our reanalysis, in the “alive” task, 1200 trials (200 trials for each category) were divided into 10 sequences, each containing 120 trials, with 20 trials from each category. Each trial consisted of a fixation cross for a random duration between 500 ms and 1000 ms, followed by an image in the center of the screen for 100 ms. Participants had 1000 ms to judge whether the image depicted a living object. A response was confirmed by the fixation cross filling in; otherwise, the screen displayed “Too late!” (Fig. 2E). Similarly, in the passive viewing task, 3600 trials (600 trials per category) were divided into 10 sequences of 360 trials, each containing 60 trials per category. Each trial consisted of a 100 ms image presentation followed by a 50 ms inter-stimulus interval. Participants were instructed to monitor a fixation bullseye and respond by pressing a button whenever it changed color to red (Fig. 2D). For further procedural details, see Shatek et al.^24^.

### 2.5 EEG data analysis

#### 2.5.1 EEG preprocessing

Preprocessed EEG data were obtained directly from the original datasets. As described in the source publication^23,24^, signals had been re-referenced to the average reference, low-pass filtered at 100 Hz, high-pass filtered at 0.1 Hz, and down-sampled to 250 Hz. Epochs were constructed from 100 ms prior to 600 ms after each image presentation for Experiment 1, and from 300 ms before each stimulus appeared on the screen to 1000 ms after stimulus onset for Experiment 2. No further preprocessing steps were applied.

#### 2.5.2 Stimulus decoding

To examine the effect of electrode density on decoding performance, we performed time-resolved multivariate pattern analysis (MVPA)^29^ on preprocessed EEG data using CoSMoMVPA^30^ in MATLAB. Neural responses were decoded separately for five electrode density levels (16, 32, 64, 96, and 128 electrodes), obtained by sampling subsets from the original 128-channel montage (Fig. 1; Supplementary Table 1). Standard electrode configurations commonly used in low-density EEG systems were followed, ensuring that subsets were evenly distributed across the scalp.

In Experiment 1, for each electrode density level, we decoded EEG epochs elicited by grating stimuli varying across four visual features: orientation, contrast, spatial frequency, and color (each with four levels). In Experiment 2, we decoded six natural image categories: living animals (whales), living plants (flowers), non-living moving artificial objects (trains), non-living moving natural objects (waterfalls), non-living still artificial objects (cups), and non-living still natural objects (rocks). At each time point of each epoch, voltage values across all electrodes within a given subset were concatenated into a feature vector, such that each sample corresponded to the multivariate scalp pattern of one trial at one time point. Regularized linear discriminant analysis (LDA) classifiers were trained and tested using a leave-one-sequence-out cross-validation procedure. For each fold, classifiers were trained on data from all but one sequence and tested on the held-out sequence, cycling through all sequences. Classification was performed separately for each visual feature at each stimulus presentation rate in Experiment 1, and for each task in Experiment 2, across all time points and electrode density levels. Decoding accuracy, defined as the proportion of correctly classified trials, was assessed for each combination of time point, feature or category, and electrode density, and compared against theoretical chance levels (0.25 for grating features; 0.167 for natural image categories). To summarize decoding performance, we quantified decoding strength for each participant as the mean decoding accuracy within the time interval identified as significant by Bayes factors (BF) > 10 (see Section 2.5.3).

To obtain an estimation of spatial contributions to decoding performance, we performed a channel-searchlight analysis. For each EEG channel, we selected the nearest 4 neighbouring channels and performed the same decoding analysis as above, storing the accuracy at the centre channel, obtaining a time-varying spatial map of cross-validated decoding accuracies for each participant.

#### 2.5.3 Statistical inference

To statistically assess whether group-level decoding accuracies exceeded chance, we computed Bayes factors (BF) ^31–33^ using the Jeffreys–Zellner–Siow (JZS) prior with a scale factor of 0.707, as implemented in the BayesFactor R package^34^ and its corresponding implementation for time-series neuroimaging data^33^. The null hypothesis was defined as decoding at chance level (0.25 in Experiment 1; 0.167 in Experiment 2), and the alternative hypothesis prior was an interval ranging from small effect sizes to infinity, accounting for small above-chance results as a result of noise^31,33^. A BF quantifies the relative evidence for the alternative compared to the null, with BF > 10 interpreted as strong evidence for above-chance decoding following established conventions^35,36^.

The onset of above-chance decoding was defined as the first of three consecutive time points with BF > 10. To be able to compare onset and peak times, we calculated 95% confidence intervals by using a leave-two-participants-out jackknifing approach^37,38^, where we calculated the onset and peak for all possible leave-two-out permutations (n = 120 for 16 participants in Experiment 1 and n = 276 for 24 participants in Experiment 2), and took the 95th percentile of the resulting distributions.

To assess the effect of electrode density on decoding strength, we fitted linear mixed-effects models (LMMs) with electrode density as a continuous predictor and participant as a random intercept.

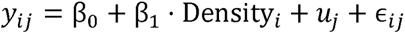

where *y_ij_* denotes the decoding strength for participant *j* at density level *i*, *β*_0_ is the intercept, *β*_1_ is the fixed effect of electrode density, *u_j_* is the participant-specific random intercept, and *ε_ij_* is the residual error term.

## 3. Results

To address how electrode density influences decoding performance, we systematically examined decoding accuracy across five electrode configurations (16, 32, 64, 96, and 128 channels) while decoding four levels of low-level visual grating features (orientation, contrast, spatial frequency, and color) in Experiment 1, and high-level natural image categories spanning six semantic classes in Experiment 2.

### 3.1 Dynamic visual grating feature coding

Using multivariate pattern analysis (MVPA), we decoded feature-specific information for orientation, contrast, spatial frequency, and color at 6.67 Hz across electrode densities from 16 to 128 channels (Fig. 3A-D, top panel). Bayes factors (BF), shown below the head maps (Fig. 3A-D, middle panel), provided strong evidence for above-chance decoding (BF > 10) across all features and electrode configurations, indicating that even the sparsest 16-channel montage was sufficient to capture feature-specific neural signals. To evaluate decoding sensitivity across electrode configurations, decoding strength was quantified for each participant as the mean decoding accuracy within the BF-defined significant interval (BF > 10). Linear mixed-effects models, with electrode density as a continuous predictor and participant as a random intercept, revealed robust positive effects of electrode density on decoding strength for orientation (β = 2.34 × 10□□, p < 0.001), spatial frequency (β = 7.80 × 10□□, p < 0.001), color (β = 2.84 × 10□□, p < 0.001), and contrast (β = 7.10 × 10□□, p < 0.001). These results indicated that decoding strength increased systematically with electrode density across all stimulus features, yet all effects were observed on top of uniformly above-chance decoding across densities.

**Fig. 3.**
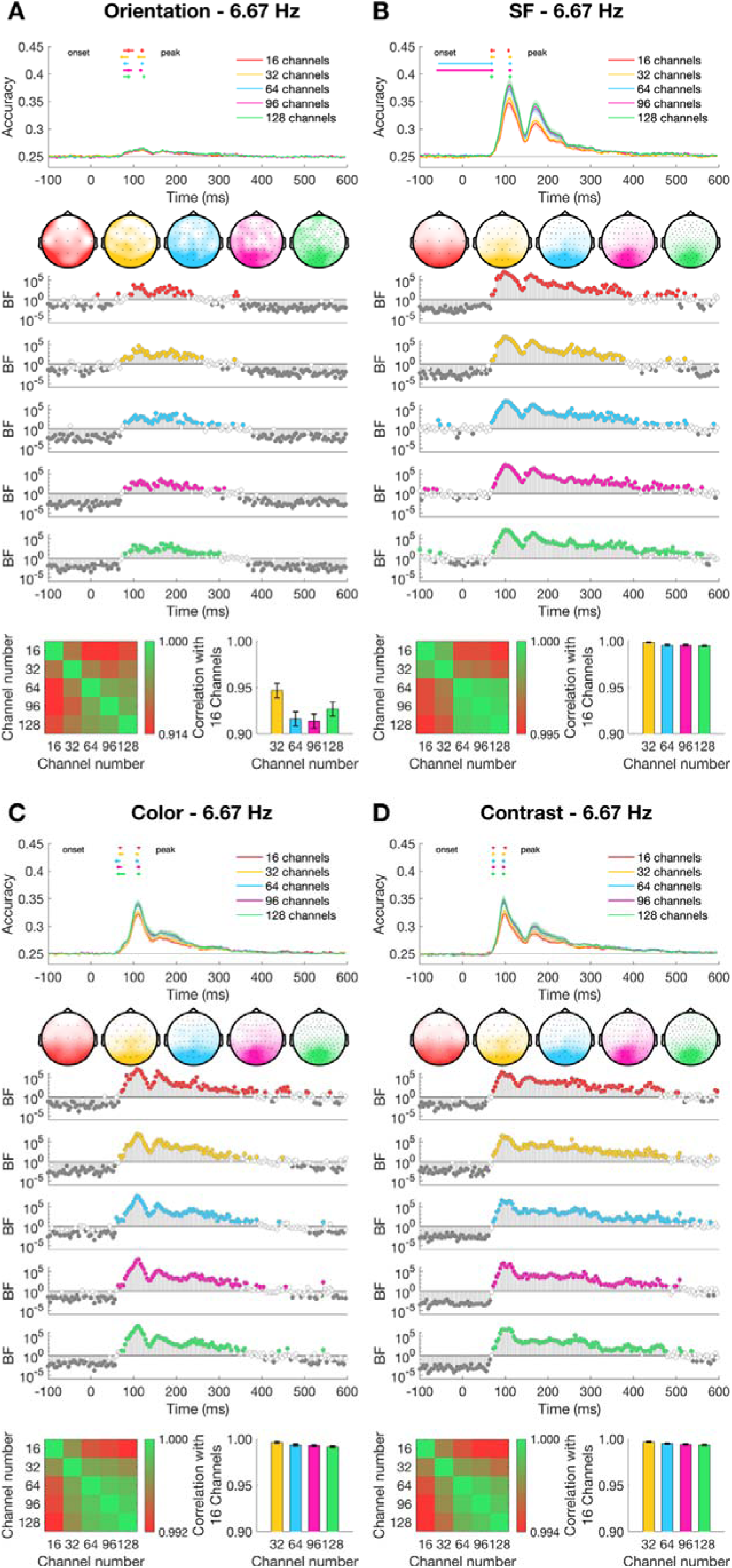
Dynamics of visual coding for orientation, spatial frequency, color, and contrast at a 6.67 Hz stimulus presentation rate with varying electrode density. (A) The time course of decoding accuracy for orientation with varying electrode density at a 6.67 Hz presentation rate. Confidence intervals for the onsets and peaks of individual electrode densities are plotted above the decoding traces. The head maps illustrate the channel clusters with the highest feature information at the peak of decoding, based on results from a channel searchlight analysis. Bayes Factors for classification evidence compared to chance (0.25) are plotted below: grey denotes evidence for the null (BF < 0.1), white denotes inconclusive evidence (0.1 ≤ BF ≤ 10), and colored denotes strong evidence for above-chance decoding (BF > 10). In the bottom row, the correlation coefficient matrix across different electrode densities and the corresponding correlation coefficient bar plot are displayed. (B–D) Same as (A), but for spatial frequency (SF; B), color (C), and contrast (D).

We next examined temporal consistency across electrode densities. Group-level decoding onset, defined as the earliest time point with three consecutive samples exceeding a Bayes factor of 10, consistently emerged rapidly after stimulus onset, with peak latencies clustering around 100–120 ms across all electrode configurations (Supplementary Table 2). Moreover, pairwise correlations of decoding time courses across densities revealed extremely high consistency, with the minimum correlation between the 16-channel montage and higher-density configurations exceeding r = 0.916 (orientation: 0.916; spatial frequency: 0.995; color: 0.992; contrast: 0.994; all p < 0.001; Fig. 3A-D, bottom panel). Finally, spatial consistency was assessed using topographical decoding maps. For all features, decoding-related activity localized reliably to occipital and parietal regions, with substantial spatial overlap across electrode densities (Fig. 3A-D, head maps), indicating stable spatial topographies of feature-specific processing.

Together, these findings demonstrate that, despite systematic differences across electrode densities, accurate, temporally consistent, and spatially overlapping decoding of multiple visual grating features remains robust, underscoring the efficiency of low-density EEG for neural decoding.

Next, we examined whether these findings generalize to a faster presentation rate of 20.00 Hz. Applying the same multivariate pattern analyses to stimuli presented at 20.00 Hz, we again observed robust above-chance decoding (BF > 10) for all four features across electrode densities, confirming that even sparse montages retained reliable feature selectivity (Fig. 4A-D, top and middle panels). Linear mixed-effects models revealed positive effects of electrode density for all four features. The effects were robust for spatial frequency (β = 1.35 × 10□□, p < 0.001), color (β = 1.04 × 10□□, p < 0.001), and contrast (β = 3.36 × 10□□, p = 0.007), while orientation showed a weaker but consistent trend in the same direction (β = 1.10 × 10□□, p = 0.058). These results indicated that decoding strength systematically increased with electrode density, while all four features remained robustly decodable.

**Fig. 4.**
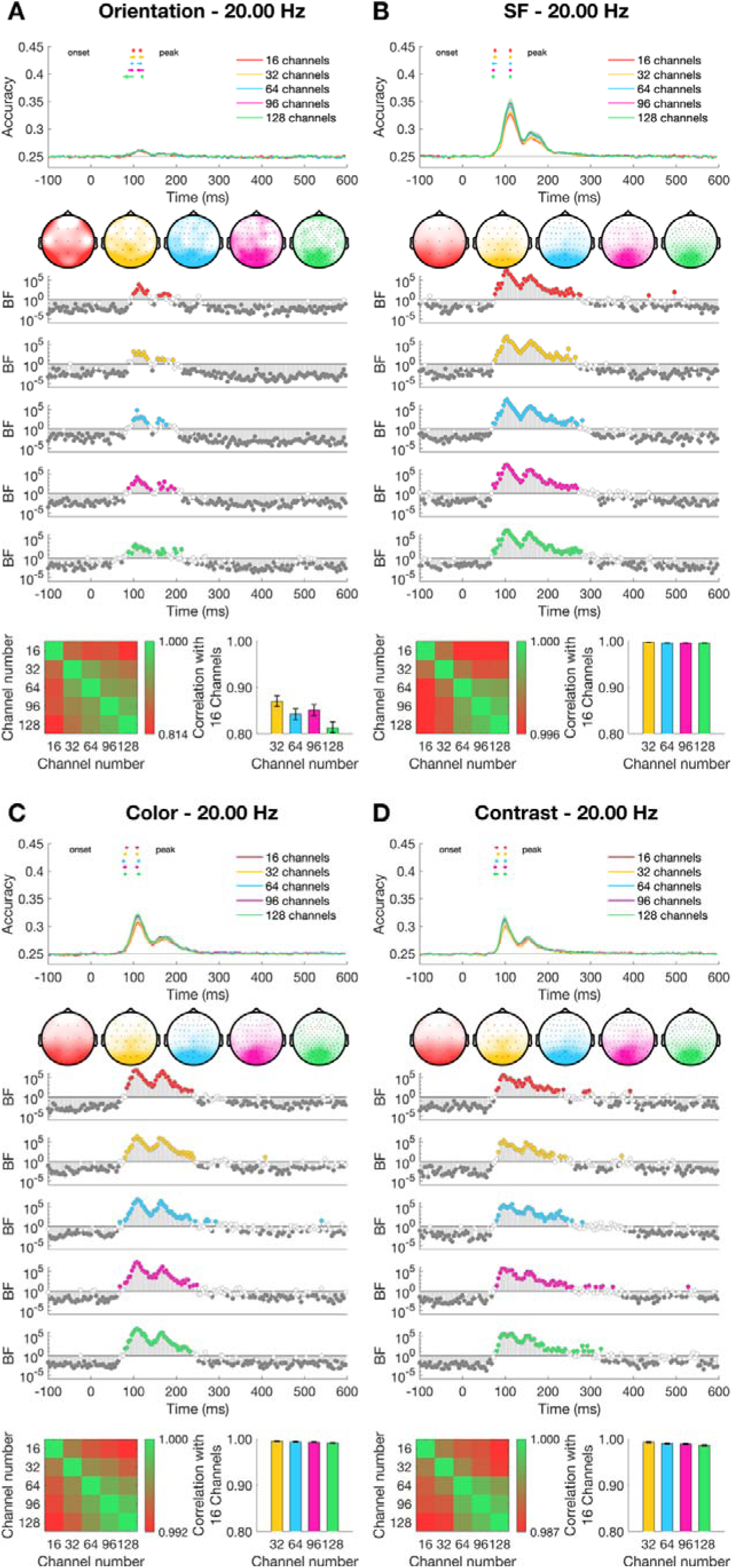
Dynamics of visual coding for orientation, spatial frequency, color, and contrast at a 20.00 Hz stimulus presentation rate with varying electrode density. (A) The time course of decoding accuracy for orientation with varying electrode density at a 20.00 Hz presentation rate. Confidence intervals for the onsets and peaks of individual electrode densities are plotted above the decoding traces. The head maps illustrate the channel clusters with the highest feature information at the peak of decoding, based on results from a channel searchlight analysis. Bayes Factors for classification evidence compared to chance (0.25) are plotted below: grey denotes evidence for the null (BF < 0.1), white denotes inconclusive evidence (0.1 ≤ BF ≤ 10), and colored denotes strong evidence for above-chance decoding (BF > 10). In the bottom row, the correlation coefficient matrix across different electrode densities and the corresponding correlation coefficient bar plot are displayed. (B–D) Same as (A), but for spatial frequency (SF; B), color (C), and contrast (D).

Temporal dynamics were preserved across electrode densities. At the group level, decoding onsets consistently occurred rapidly and peaks clustered around 96–124 ms across all densities (Supplementary Table 3). Temporal profile similarity remained high, with pairwise correlations between the 16-channel montage and higher-density configurations exceeding r = 0.814 for orientation, 0.996 for spatial frequency, 0.992 for color, and 0.987 for contrast (all p < 0.001; Fig. 4A-D, bottom panel). Finally, topographical decoding maps again localized feature-selective activity to occipital–parietal regions with substantial overlap across densities (Fig. 4A-D, head maps).

Together, these findings demonstrate that decoding performance remains robust across electrode densities under both moderate (6.67 Hz) and accelerated (20.00 Hz) stimulation, supporting the reliability and efficiency of low-density EEG for feature-specific neural decoding.

### 3.2 Dynamic natural stimuli coding

We next examined whether the decoding consistency across electrode densities extends to high-level, naturalistic visual input. To test this, we applied the same multivariate pattern analysis to EEG data collected during the viewing of natural images from six semantic categories: living animals (whales), living plants (flowers), non-living moving artificial objects (trains), non-living moving natural objects (waterfalls), non-living still artificial objects (cups), and non-living still natural objects (rocks). Analyses were conducted separately for the categorization task (Fig. 5A) and the passive viewing task (Fig. 5B), across five electrode densities (16, 32, 64, 96, and 128 channels).

**Figure 5.**
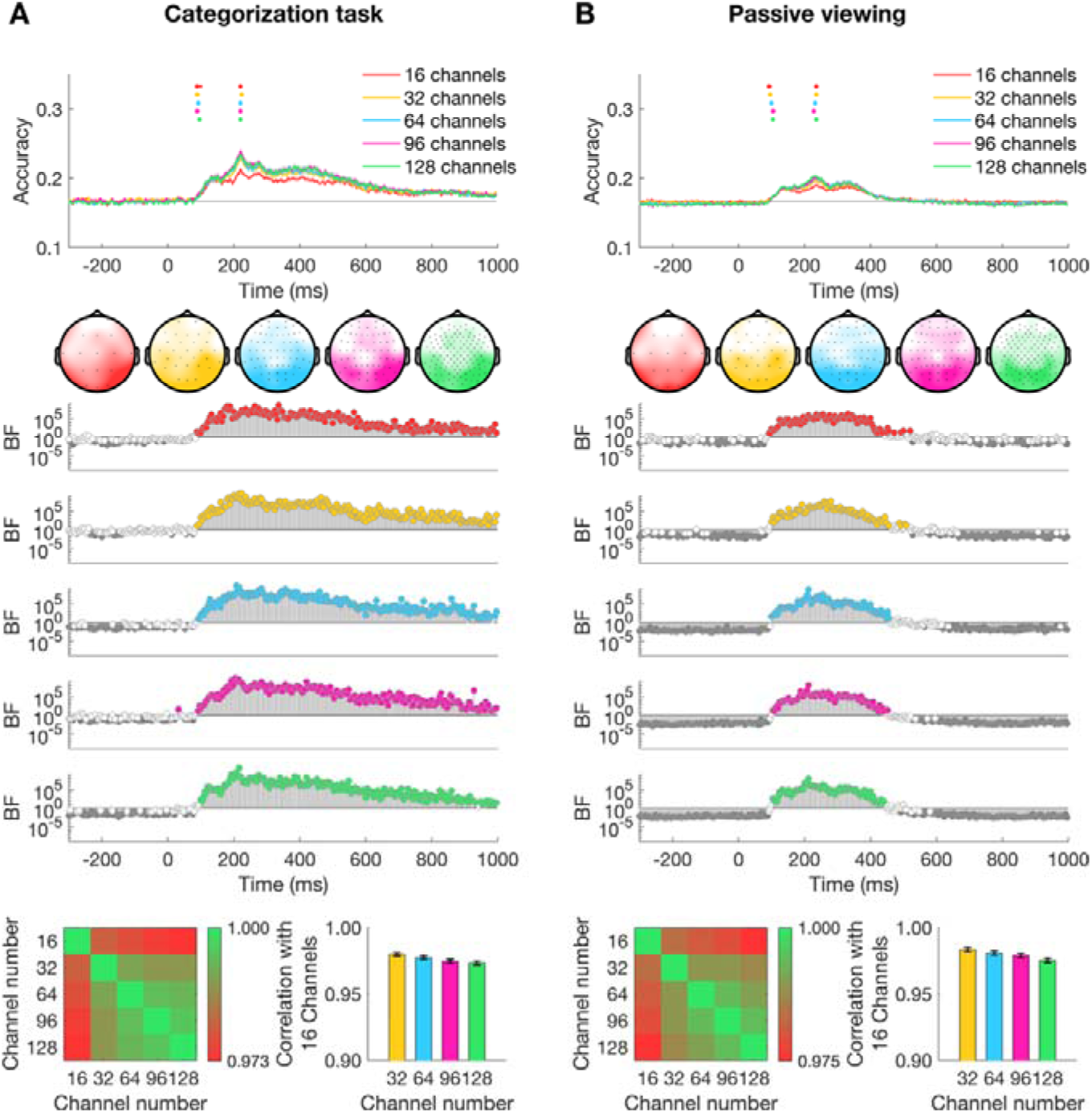
Dynamics of visual coding for natural stimuli at categorization and passive viewing tasks with varying electrode density. (A) The time course of decoding accuracy at categorization task with varying electrode density. Confidence intervals for the onsets and peaks of individual electrode densities are plotted above the decoding traces. The head maps illustrate the channel clusters with the highest categorical information at the peak of decoding, based on results from a channel searchlight analysis. Bayes Factors for classification evidence compared to chance (0.167) are plotted below: grey denotes evidence for the null (BF < 0.1), white denotes inconclusive evidence (0.1 ≤ BF ≤ 10), and colored denotes strong evidence for above-chance decoding (BF > 10). In the bottom row, the correlation coefficient matrix across different electrode densities and the corresponding correlation coefficient bar plot are displayed. (B) Same as (A), but at passive viewing task.

For the categorization task (Fig. 5A), decoding accuracy for object category was reliably above chance (BF > 10) across all electrode densities (Fig. 5A, top and middle panels). To assess how electrode density modulated sensitivity, we quantified decoding strength for each participant as the mean accuracy within the BF-defined significant interval. Linear mixed-effects modelling, treating electrode density as a continuous predictor, revealed a clear positive association between density and decoding strength (β = 5.59 × 10□□, p < 0.001). Despite this density-dependent gain, the temporal dynamics were highly consistent across montages: decoding emerged rapidly post-stimulus and peaked between ∼220 ms (Supplementary Table 4). In addition, temporal profile similarity was extremely high, with pairwise correlations between the 16-channel montage and higher-density configurations exceeding r = 0.973 (p < 0.001; Fig. 5A, bottom panel). Finally, topographical decoding maps localized categorical information to occipital–parietal regions, showing substantial spatial overlap across densities (Fig. 5A, head maps) and indicating a robust spatiotemporal signature of categorical processing.

For the passive viewing task (Fig. 5B), decoding accuracy was again reliably above chance across all electrode densities. Linear mixed-effects models likewise revealed a significant positive effect of electrode density on decoding strength (β = 6.47 × 10□□, SE = 9.28 × 10□□, t(118) = 6.97, p < 0.001). Time courses were broadly similar across montages, with decoding rising rapidly after stimulus onset and peaking around ∼230 ms (Fig. 5B, top and middle panels; Supplementary Table 4). Pairwise correlations confirmed high temporal consistency, with the minimum correlation between the 16-channel montage and higher-density configurations exceeding r = 0.975 (p < 0.001; Fig. 5B, bottom panel). Spatial decoding maps again revealed occipital–parietal localization with strong overlap across electrode densities (Fig. 5B, head maps), demonstrating a stable spatial topography of category-related activity.

Together, these findings demonstrate that high-level visual information can be accurately and consistently decoded from EEG signals even with sparse electrode configurations. While denser montages confer advantages in decoding performance, the overall robustness across densities highlights the utility of low-density EEG for decoding complex, naturalistic stimuli.

## Discussion

In this study, we systematically examined how varying EEG electrode densities influence neural decoding performance for both low-level (Experiment 1) and high-level (Experiment 2) visual stimuli. As expected, decoding performance increased with electrode density. Remarkably, however, temporal decoding accuracy remained above chance even with as few as 16 electrodes, irrespective of stimulus complexity or task demands. Moreover, spatial patterns of decoding-related activity were broadly consistent across densities, indicating a degree of spatial robustness. Together, these findings underscore both the benefits of higher-density recordings and the reliability and generalizability of EEG-based neural decoding under sparse spatial sampling conditions.

Prior studies have suggested that increasing EEG electrode density enhances decoding accuracy, particularly for fine-grained perceptual tasks^10–12^, a pattern consistent with our findings. At the same time, recent studies suggest that reliable decoding can still be achieved with fewer electrodes^8,9,13,39^. However, these studies were limited in scope, focusing on dense grids over the occipital cortex^8^, resting-state functional connectivity^13^, or motor imagery BCIs^9, 39^, rather than systematically varying electrode density across the whole brain in visual paradigms. To our knowledge, the present study is the first to directly compare neural decoding across multiple standard electrode configurations within the same visual tasks. By examining both low- and high-level visual stimuli, our results extend previous findings and provide systematic evidence that decoding accuracy and temporal dynamics remain robust even under sparse spatial sampling.

Although task-state EEG signals originate from anatomically localized sources, such as early visual areas in the occipital cortex involved in encoding low-level features^14,40^, they give rise to scalp potentials that are both locally enhanced and spatially distributed due to volume conduction^15–18^. Specifically, neural activity from focal sources propagates through the brain and skull, producing widespread scalp fields that extend well beyond the cortical origin. At the same time, these fields tend to exhibit maximal amplitude near their generating sources, which explains the consistently observed topography maps over occipital regions in Experiment 1. This dual property of EEG, global spread with regional specificity, enables sparse but widely distributed electrode arrays to capture the key components of multivariate signal patterns^8,9,13^. As long as the electrode montage ensures full-head coverage, even low-density configurations can retain sufficient information for effective decoding. Our findings support this principle: despite substantial reductions in electrode count, we observed reliable decoding of both temporal dynamics and spatial structures, across both low- and high-level visual tasks.

The visual tasks in this study employed rapid serial visual presentation (RSVP) paradigms, which are widely used to approximate the continuous nature of visual processing^41–46^. A potential concern is that this design may introduce overlap between consecutive epochs. However, all analyses were strictly time-locked to stimulus onset, and any overlapping activity would be equally distributed across conditions because trials were randomized. Thus, potential contributions from subsequent stimuli are unlikely to systematically bias the reported decoding accuracy. In addition, we also found several patterns in visual decoding performance that echo previous studies^23,24^. Firstly, in Experiment 1, orientation decoding was relatively low (though reliable) compared to other visual features (Fig. 3 & 4) no matter in which electrode density level. Similar results have been reported previously, where EEG shows weaker orientation decoding while MEG reveals stronger signals^47,48^. Although our design was highly powered (2,560 events per orientation angle), the exclusion of cardinal and oblique orientations likely reduced sensitivity^48^. Moreover, features such as color, contrast, and spatial frequency evoke stronger and more distributed responses (e.g., high contrast drives robust signals^49^), which are easier to detect at the scalp. In contrast, orientation information may be downweighted at higher levels of visual processing because robust object perception requires rotational invariance^50,51^. This may explain why orientation decoding is weaker relative to other features, despite its importance for local edge and shape detection. Secondly, decoding accuracy was lower in the 20.00 Hz (Fig. 4) compared to the 6.67 Hz (Fig. 3) condition, despite overall similar temporal dynamics. We interpret this reduction as arising from decreased signal-to-noise ratio at higher presentation rates. A similar pattern was observed in Experiment 2, where decoding accuracy was lower in the passive viewing than in the categorization task, which can likewise be attributed to reduced task engagement and thus lower signal-to-noise ratio. Thirdly, low-level feature coding exhibited a pronounced double-peak response (Fig. 3 &4), with an early peak around 90–130 ms and a later peak around 160–200 ms corresponding to the P100 and N1/N170 components. These likely reflect distinct feedforward and feedback stages of visual processing, consistent with previous EEG decoding studies^4,6,50^. In contrast, high-level tasks (Fig. 5) such as categorization and passive viewing showed more sustained decoding with later onset and latency. This pattern suggests that while early visual features are processed rapidly through feedforward mechanisms, higher-level representations may depend more on recurrent processing and feedback, supporting prolonged and temporally extended decoding^23,24^. Importantly, these neurophysiological trends were robust across electrode density levels, supporting the broader conclusion that reliable visual decoding of both low- and high-level processes can be achieved even with reduced electrode coverage.

A key strength of this study lies in the systematic manipulation of electrode density across two distinct experimental datasets, which provides robust validation of the findings. While this approach provides robust validation under realistic recording conditions, it does not address the potential benefits of optimizing electrode placement for individual participants or specific tasks. Future studies could explore individualized electrode selection strategies, such as identifying the most informative channels based on subject-specific signal-to-noise ratios, decoding performance, or task-related activation patterns^8,39^, to further enhance the efficiency of low-density EEG systems. Another promising direction for future work is to investigate whether decoding based on narrowband signals, such as alpha power, yields comparable results to broadband approaches. Numerous studies suggest that alpha-band signals can carry robust information about visual stimuli and attention, particularly under conditions that allow for sustained neural engagement^52, 53^. Alpha oscillations and their lateralization can be reliably induced by visual stimulation^54^, and have also been shown to be evoked by novel visual stimuli^55, 56^. Therefore, future studies employing longer stimulus durations or tasks requiring sustained attention could directly test the added value of frequency-specific features in decoding performance.

Our results demonstrated that reliable decoding can be achieved with fewer electrodes, which supported the broadly distributed activities across the whole brain and opened new possibilities for scalable applications such as mobile BCIs, clinical monitoring, and real-world cognitive assessment^46, 57–59^. These findings carry important implications for the development of portable and cost-effective EEG systems.

## Supporting information

supplemental information_clean

## Data and Code Availability

This study utilized publicly available datasets from OpenNeuro: Experiment 1 (https://doi.org/10.18112/openneuro.ds004357.v1.0.1) and Experiment 2 (https://doi.org/10.18112/openneuro.ds003885.v1.0.7). Code is publicly available at https://osf.io/xu2he/

## Author contributions

Conceptualization, L.H., M.V., T.G.; Methodology, L.H., T. G.; Formal Analysis, L.H.; Visualization L.H.; Writing – Original Draft, L. H.; Writing – Review and Editing, L. H., M.V., T. G.

## Ethics Statement

This work used publicly available datasets that were originally collected under the University of Sydney Human Research Ethics Committee approvals. No additional data collection or ethics approval was required for the present analyses.

