## supplemental information_clean for "Robust Neural Decoding with low-density EEG"

1. **Supplementary Table 1**. Electrode labels for each electrode density configuration.
2. **Supplementary Table 2**. Decoding performance (onset time, peak time, maximum decoding accuracy, and decoding strength) for orientation, contrast, spatial frequency, and color at 6.67 Hz for each electrode density configuration.
3. **Supplementary Table 3**. Decoding performance (onset time, peak time, maximum decoding accuracy, and decoding strength) for orientation, contrast, spatial frequency, and color at 20.00 Hz for each electrode density configuration.
4. **Supplementary Table 4**. Decoding performance (onset time, peak time, maximum decoding accuracy, and decoding strength) for categorization and passive viewing tasks for each electrode density configuration.

**Supplementary Table 1**. Electrode labels for each electrode density configuration. Each lower-density montage was constructed as a spatially balanced subset of the original 128-channel configuration.

| **Electrode Density** | **Number of Channels** | **Electrode Labels** |
| --- | --- | --- |
| 16-channel | 16 | Fp1, Fz, F3, C3, T7, Pz, P3, O1, Oz, O2, P4, Cz, C4, T8, F4, Fp2 |
| 32-channel | 32 | *(16-channel set)* + F7, FT9, FC5, FC1, TP9, CP5, CP1, P7, P8, TP10, CP6, CP2, FT10, FC6, FC2, F8 |
| 64-channel | 64 | *(32-channel set)* + AF7, AF3, AFz, F1, F5, FT7, FC3, C1, C5, TP7, CP3, P1, P5, PO7, PO3, POz, PO4, PO8, P6, P2, CPz, CP4, TP8, C6, C2, FC4, FT8, F6, AF8, AF4, F2 |
| 96-channel | 96 | *(64-channel set)* + F9, AFF1h, FFC1h, FFC5h, FTT7h, FCC3h, CCP1h, CCP5h, TPP7h, P9, PPO9h, PO9, O9, OI1h, PPO1h, CPP3h, CPP4h, PPO2h, OI2h, O10, PO10, PPO10h, P10, TPP8h, CCP6h, CCP2h, FCC4h, FTT8h, FFC6h, FFC2h, AFF2h, F10 |
| 128-channel | 128 | *(96-channel set)* + AFp1, AFF5h, FFT9h, FFT7h, FFC3h, FCC1h, FCC5h, FTT9h, TTP7h, CCP3h, CPP1h, CPP5h, TPP9h, POO9h, PPO5h, POO1, POO2, PPO6h, POO10h, TPP10h, CPP6h, CPP2h, CCP4h, TTP8h, FTT10h, FCC6h, FCC2h, FFC4h, FFT8h, FFT10h, AFF6h, AFp2 |

**Supplementary Table 2**. Decoding performance (onset time, peak time, maximum decoding accuracy, and decoding strength) for orientation, contrast, spatial frequency (SF), and color at 6.67 Hz for each electrode density configuration.

| Feature | Electrode Density | Onset  Time (ms) | Peak  Time (ms) | Max Accuracy | Decoding  Strength |
| --- | --- | --- | --- | --- | --- |
| orientation | 16 | 100 | 116 | 0.259 | 0.255 |
| orientation | 32 | 100 | 116 | 0.260 | 0.255 |
| orientation | 64 | 96 | 116 | 0.261 | 0.256 |
| orientation | 96 | 96 | 112 | 0.262 | 0.256 |
| orientation | 128 | 84 | 120 | 0.263 | 0.256 |
| SF | 16 | 76 | 112 | 0.326 | 0.262 |
| SF | 32 | 76 | 112 | 0.330 | 0.276 |
| SF | 64 | 72 | 112 | 0.341 | 0.278 |
| SF | 96 | 72 | 112 | 0.346 | 0.281 |
| SF | 128 | 72 | 112 | 0.348 | 0.281 |
| Color | 16 | 84 | 108 | 0.307 | 0.273 |
| Color | 32 | 80 | 108 | 0.309 | 0.262 |
| Color | 64 | 76 | 112 | 0.317 | 0.261 |
| Color | 96 | 80 | 108 | 0.319 | 0.277 |
| Color | 128 | 80 | 112 | 0.321 | 0.279 |
| Contrast | 16 | 80 | 100 | 0.300 | 0.260 |
| Contrast | 32 | 84 | 100 | 0.302 | 0.260 |
| Contrast | 64 | 80 | 100 | 0.311 | 0.268 |
| Contrast | 96 | 76 | 100 | 0.312 | 0.259 |
| Contrast | 128 | 76 | 100 | 0.314 | 0.266 |

**Supplementary Table 3**. Decoding performance (onset time, peak time, maximum decoding accuracy, and decoding strength) for orientation, contrast, spatial frequency (SF), and color at 20.00 Hz for each electrode density configuration.

| Feature | Electrode Density | Onset  Time (ms) | Peak  Time (ms) | Max Accuracy | Decoding  Strength |
| --- | --- | --- | --- | --- | --- |
| orientation | 16 | 88 | 120 | 0.26156 | 0.25523 |
| orientation | 32 | 72 | 112 | 0.26197 | 0.25598 |
| orientation | 64 | 80 | 120 | 0.26468 | 0.25656 |
| orientation | 96 | 88 | 116 | 0.26489 | 0.25718 |
| orientation | 128 | 88 | 124 | 0.26576 | 0.25804 |
| SF | 16 | 68 | 108 | 0.34681 | 0.27057 |
| SF | 32 | 68 | 112 | 0.35592 | 0.27268 |
| SF | 64 | 68 | 112 | 0.37242 | 0.27544 |
| SF | 96 | 68 | 112 | 0.37819 | 0.27817 |
| SF | 128 | 68 | 112 | 0.38101 | 0.27937 |
| Color | 16 | 68 | 112 | 0.32209 | 0.26325 |
| Color | 32 | 68 | 108 | 0.32576 | 0.26712 |
| Color | 64 | 60 | 108 | 0.33693 | 0.26699 |
| Color | 96 | 64 | 112 | 0.34081 | 0.26762 |
| Color | 128 | 64 | 112 | 0.34248 | 0.26762 |
| Contrast | 16 | 72 | 100 | 0.32445 | 0.26721 |
| Contrast | 32 | 72 | 96 | 0.32843 | 0.26571 |
| Contrast | 64 | 72 | 96 | 0.34254 | 0.27036 |
| Contrast | 96 | 72 | 96 | 0.34744 | 0.27338 |
| Contrast | 128 | 72 | 96 | 0.34959 | 0.27247 |

**Supplementary Table 4**. Decoding performance (onset time, peak time, maximum decoding accuracy, and decoding strength) for categorization and passive viewing tasks for each electrode density configuration.

| Task | Electrode Density | Onset  Time (ms) | Peak  Time (ms) | Max Accuracy | Decoding  Strength |
| --- | --- | --- | --- | --- | --- |
| categorization | 16 | 88 | 220 | 0.21365 | 0.18853 |
| categorization | 32 | 88 | 224 | 0.22931 | 0.19371 |
| categorization | 64 | 92 | 220 | 0.23358 | 0.19495 |
| categorization | 96 | 88 | 220 | 0.23944 | 0.19669 |
| categorization | 128 | 96 | 220 | 0.23368 | 0.19572 |
| Passive view | 16 | 92 | 236 | 0.19044 | 0.1799 |
| Passive view | 32 | 96 | 236 | 0.19689 | 0.18261 |
| Passive view | 64 | 100 | 232 | 0.20042 | 0.18673 |
| Passive view | 96 | 104 | 228 | 0.20112 | 0.18655 |
| Passive view | 128 | 104 | 236 | 0.2035 | 0.18762 |
